# A role for the Erk MAPK pathway in modulating SAX-7/L1CAM-dependent locomotion in *Caenorhabditis elegans*

**DOI:** 10.1101/2021.03.09.434630

**Authors:** Melinda Moseley-Alldredge, Seema Sheoran, Hayoung Yoo, Calvin O’Keefe, Janet E. Richmond, Lihsia Chen

## Abstract

L1CAMs are immunoglobulin cell adhesion molecules that play important roles in the development and function of the nervous system. In addition to being associated with autism and schizophrenia spectrum disorders, impaired L1CAM function also underlies the X-linked L1 syndrome, which encompasses a group of neurological conditions, including spastic paraplegia and congenital hydrocephalus. Previous studies on both vertebrate and invertebrate L1CAMs established conserved roles that include axon guidance, dendrite morphogenesis, synapse development, and maintenance of neural architecture. We previously identified a genetic interaction between the *C. elegans* L1CAM encoded by the *sax-7* gene and RAB-3, a GTPase that functions in synaptic neurotransmission; *rab-3; sax-7* animals exhibit synthetic locomotion abnormalities and neuronal dysfunction. In this study, we examine the significance of this genetic interaction and show that this synergism also occurs when loss of SAX-7 is combined with mutants of other genes encoding key players of the synaptic vesicle cycle. In contrast, *sax-7* does not interact with genes that function in synaptogenesis. These findings suggest a post-developmental role for *sax-7* in the regulation of synaptic activity. To further assess this possibility, we conducted electrophysiological recordings and ultrastructural analyses at neuromuscular junctions. Lastly, we performed a forward genetic screen for suppressors of the *rab-3; sax-7* synthetic phenotypes, uncovering a role for the Mitogen-activated Protein Kinase (MAPK) pathway in promoting coordinated locomotion.

## Introduction

The immunoglobulin superfamily of cell adhesion molecules (IgCAMs) is essential in the development and function of the nervous system, playing roles that range from axon guidance and fasciculation to synapse formation and regulation of synaptic activity. Single-pass transmembrane L1CAMs are a well-characterized sub-family of neuronal IgCAMs that are highly conserved in vertebrates, *Drosophila*, and *C. elegans*. Studies have established conserved L1CAM roles in axon guidance, myelination, and fasciculation, as well as dendrite morphogenesis and synaptogenesis (Chen and Zhou 2010; Hortsch *et al.* 2014; Sundararajan *et al.* 2019). Consistent with these neurodevelopmental roles, mutations in human L1CAM-encoding genes, L1, NrCAM, CHL1, and neurofascin, are strongly implicated in a variety of neurological disorders. For example, mutations in the L1 gene are directly linked to the congenital X-linked L1 syndrome that encompass a range of neurological conditions, including hydrocephalus, spastic paraplegia, intellectual disability, and corpus callosum hypoplasia (Fransen *et al.* 1994; Vits *et al.* 1994; Fransen *et al.* 1995; Van Camp *et al.* 1996). Interestingly, the severity or manifestation of these conditions can vary even among family members carrying the same L1 mutation, suggesting the presence of other genetic variants that can interact with L1 to modify disease expressivity or penetrance (Fryns *et al.* 1991; Jouet *et al.* 1995; Schrander-Stumpel *et al.* 1995). This variability, particularly with respect to hydrocephalus, was also observed in independently generated L1 knockout mice (Dahme *et al.* 1997; Fransen *et al.* 1998). Consistent with the notion that genetic interactions with L1 underlie disease severity, a subsequent study in mice uncovered genetic modifier loci that influenced L1-associated hydrocephalus (Tapanes-Castillo *et al.* 2010).

In addition to developmental disorders, both genome-wide association studies and individual patient analyses have also implicated L1CAMs in specific behavioral disorders. NrCAM is associated with autism spectrum disorder and susceptibility to addiction; CHL1 linked to schizophrenia (Sakurai *et al.* 2002; Ishiguro *et al.* 2006; Sakurai *et al.* 2006; Tam *et al.* 2010; Ayalew *et al.* 2012; Shaltout *et al.* 2013; Zhong *et al.* 2015). Similarly, L1, NrCAM, and CHL1 knockout mice exhibit altered social and exploratory behaviors (Fransen *et al.* 1998; Law *et al.* 2003; Moy *et al.* 2009). As most genetic mutations implicated in behavioral disorders have mild effects on behavior individually, it has been challenging to determine how these genetic variants contribute to behavioral disorders. It is hypothesized that it is the genetic interactions between two or more variants that are required for the manifestation of behavioral conditions (Owen *et al.* 2016; Wamsley and Geschwind 2020). Identifying such genetic interactions will help in understanding the function of the L1CAMs associated genetic networks in behavioral outputs.

We previously identified several distinct genetic interactions with the *C. elegans* canonical L1CAM encoded by the *sax-7* gene (Opperman *et al.* 2015). Despite established roles in neural architecture maintenance, dendrite morphogenesis and organization, axon branching, and axon-dendrite fasciculation (Wang *et al.* 2005; Dong *et al.* 2013; Diaz-Balzac *et al.* 2015; Diaz-Balzac *et al.* 2016; Yip and Heiman 2018; Chen *et al.* 2019), *sax-7* null animals do not have obvious locomotory abnormalities. However, in sensitized *rab-3* or *unc-13* genetic backgrounds, *sax-7* null animals exhibit synthetic or synergistic uncoordinated (Unc) locomotion and neuronal dysfunction. Intriguingly, late-onset, transient SAX-7 expression can suppress these phenotypes but cannot rescue the neural architecture maintenance defects also exhibited in *sax-7* mutants (Opperman *et al.* 2015). These findings are consistent with a post-developmental role for sax-7 in directing coordinated locomotion. Taken together with known functions of *rab-3* and *unc-13* in exocytosis and a role for CHL1 in mammalian synaptic vesicle recycling (Nonet *et al.* 1997; Richmond *et al.* 1999; Leshchyns’ka *et al.* 2006), these findings suggest *sax-7* may have a role in synaptic transmission. In this study, we use genetics, electrophysiology, and image analyses of *C. elegans* synapses to assess how *sax-7* promotes coordinated locomotion. These results together with further genetic analyses of sax-7 function uncovered a role for the Mitogen-Activated Protein Kinase (MAPK) pathway in central cholinergic neurons to promote coordinated locomotion.

## Materials and Methods Strains

*C. elegans strains*, provided by the *Caenorhabditis* Genetics Center, were grown on nematode growth medium (NGM) plates at 21°C. N2 Bristol served as the wild-type strain (Brenner 1974). The alleles used in this study are listed by linkage groups as follows:

LGI: *unc-13*(*n2813*) (Brenner 1974)
LGII: *rab-3*(*js49*) (Nonet *et al.* 1997); *syd-1*(*ju82*) (Hallam *et al.* 2002)
LGIII: *mpk-1*(*ga117*) (Lackner *et al.* 1994)
LGIV: *sax-7*(*eq1*) (Wang *et al.* 2005)
LGV: *rpm-1*(*ju41*) (Zhen *et al.* 2000); *snb-1(e1563)* (Sandoval *et al.* 2006)
LGX: *unc-10*(*md1117*) (Koushika *et al.* 2001); *ksr-1(ok786)* (Consortium 2012)

### The strains generated in this study are as follows

LH1038: *sax-7(eq1); snb-1(e1563)*
LH1055: *rab-3(js49); sax-7(eq1); ksr-1(ok786)*
LH1079: *sax-7(eq1); unc-10(md1117)*
LH1086: *sax-7(eq22)*
LH1087: *sax-7(eq23)*
LH1098: *rab-3(js49); sax-7(eq22); tmIs1087 [P_myo-3_::Cre]*
LH1102: *rab-3(js49); sax-7(eq23)*
LH1118: *syd-1*(ju82); sax-7(*eq1*)
LH1119: *sax-7(eq1); rpm-1(ju41)*
LH1132: *rab-3(js49); sax-7(eq22); tmIs1028 [P_dpy-7_::Cre]*
LH1145: *rab-3(js49); sax-7(eq22); tmIs778 [P_rgef-1_::Cre]*
LH1190: *rab-3(js49); mpk-1(ga117)/hT2; sax-7(eq1*
LH1192: rab-3(js49); sax-7(eq22); *eqIs4 [P_rab-3_::Cre]*
LH1200: *rab-3(js49); sax-7(eq1); ksr-1(eq7)*
LH1217: *rab-3(js49); sax-7(eq22)*
LH1235: *unc-13(n2813); sax-7(eq1); ksr-1(eq7)*
LH1298*: rab-3(js49); sax-7(eq1); ksr-1(ok786); eqSi1 [P_unc17H_::ksr-1]*
LH1299*: rab-3(js49); sax-7(eq1); ksr-1(ok786); eqSi2 [P_unc17B_::ksr-1]*
LH1300*: rab-3(js49); sax-7(eq1); ksr-1(ok786); eqSi3 [P_unc17_::ksr-1]*

### Plasmids generated

pLC739: sgRNA plasmid that was used to generate the conditional *sax-7(eq22)* knock-in strain via Crispr gene-editing. The sgRNA was generated by cloning DNA oligonucleotides LC1638/39 into pRB1017, a gift from Andrew Fire (Addgene plasmid # 59936; http://n2t.net/addgene:59936; RRID:Addgene_59936) (Arribere *et al.* 2014).
pLC740: repair plasmid carrying the homology arms that was used to generate the conditional *sax-7(eq22)* knock-in strain via Crispr gene-editing. 5’ and 3’ homology arms were amplified from genomic worm DNA via PCR using primers LC1634/35 and LC1636/37 and cloned by standard restriction digestion with SpeI and Not1 into the vector carrying mCherry_loxP_myo_neoR as described (Norris *et al.* 2015).
pLC743: sgRNA plasmid carrying primers LC1671 and LC1672 that is specific to β-lactamase.
pLC755: sgRNA plasmid carrying oligonucleotides LC1778 and LC1779 that is specific to mKate2.
pLC756: sgRNA plasmid carrying oligonucleotides LC1780 and LC1781 that is specific to mKate2.
pLC759: The P*_rab-3_*::Cre-recombinase repair plasmid was generated via Gibson assembly of the *rab-3* promoter, CRE recombinase and the *tbb-2* UTR into plasmid pCFJ150-pDESTttTi5605[R4-R3], a gift from Erik Jorgensen (Addgene plasmid # 19329; http://n2t.net/addgene:19329; RRID:Addgene_19329) (Frokjaer-Jensen *et al.* 2008).

### Oligonucleotides used are listed 5’ to 3’

LC1634 ggtgactagtatcgcaaaacgaaattctcc
LC1635 ggtgactagttcgcttttcatcatcatcgct
LC1636 ggtggcggccgcatccttaacgggctccaaagcc
LC1637 ggtggcggccgccgactcgggaagaaaggag
LC1638 tcttgatgaaaagcgatcattgac
LC1639 aaacgtcaatgatcgcttttcatc
LC1671 tcttgttaatagactggatggagg
LC1672 aaaccctccatccagtctattaac
LC1778 tcttgagtcaacttcccatccaa
LC1779 aaacttggatgggaagttgactc
LC1780 tcttgcacttcaagtgcacctccga
LC1781 aaactcggaggtgcacttgaagtgc

### Generation of *sax-7(eq22)* and *sax-7(eq23)* alleles

mCherry coding sequence was added to the cytoplasmic tail of *sax-7* using CRISPR/Cas9 (Norris *et al*, 2015). The sgRNA-containing plasmid (pLC739) along with the repair plasmid (pLC740) were injected into *C. elegans* gonads as described (Norris *et al.* 2015). Worms containing an integrated P*_myo-2_*::GFP cassette were identified, and individually picked until homozygous. This strain (*eq22*) was then injected with the Cre recombinase-expressing plasmid pDD104, a gift from Bob Goldstein (Addgene plasmid # 47551; http://n2t.net/addgene:47551; RRID:Addgene_47551) (Dickinson *et al.* 2013) to excise the gene-disrupting P_myo-2_::GFP cassette, generating the *eq23* allele, which will result in SAX-7::mCherry expression.

### Tissue-specific *sax-7* knock-ins

To knock in *sax-7* expression in a tissue specific manner, we excised the gene-disrupting cassette in *sax-*7(*eq22*) using multi-copy integrated arrays expressing Cre recombinase driven by promoters of interest. For neuronal, muscle, and hypodermal expression of sax-7, we used integrated multicopy arrays (*tmIs778, tmIs1087*, and *tmIs1027*) expressing Cre recombinase that is driven by the *rgef-1, myo-3* or *dpy-7* promoters, respectively (Kage-Nakadai *et al.* 2014). We also used another neuronally-expressing Cre transgene we generated: the P*_rab-3_*::CRE. To generate this transgene, the pLC759 plasmid was injected along with coinjection marker P*_sur-5_*::GFP into *eq22* animals to generate transgenic animals carrying extrachromosomal arrays; these arrays were then integrated into the genome using a variation of a published method (Yoshina *et al.* 2016). This adapted method is as follows: the extrachromosomal arrays are first crossed into CGC62, a strain carrying a single copy insertion of P*_myo-2_*::mKate2 on LGV. Animals homozygous for P*_myo-2_*::mKate2 also carrying the extrachromosomal array were then injected with a cocktail containing two sgRNA plasmids specific to mKate2 (pLC755 and pLC756), an sgRNA plasmid specific to β-lactamase (plasmid pLC743), P*_eft-3_*::Cas9, and pCFJ90. F2 animals with positive GFP expression from the array, but negative for mKate expression were individually picked and checked for Mendelian expression of P*_sur-5_*::GFP from the array, which indicates successful targeted transgene integration (*eqIs4)*.

### Site-specific single copy transgene insertions

Single copy insertions of P*_unc-17_*::*ksr-1::gfp*, P*_unc-17H_*::*ksr-1::gfp*, and P*_unc-17B_*::*ksr-1::gfp* were generated using CRISPR/Cas9. Each transgene was inserted into the *oxTi365* MoSCI site present in the EG8082 strain (Frokjaer-Jensen *et al.* 2014). To do so, we injected each of corresponding *ksr-1-*containing MosSCI plasmids (pBC31, pBC32 and pBC33 (Coleman *et al.* 2018)) individually at 50ng/μl along, along with a cocktail containing 50ng/ul each of sgRNA-containing vectors pXW7.01 and pXW7.02 (gift of Katya Voronina, University of Montana) to cleave the universal ttTi5605 MosSCI site, and 2.5ng/μl of selection marker pCFJ90 (P*_myo-2_*::mCherry). Injected animals were incubated at 25°C for approximately 7 days, after which non-Unc, non-fluorescent F2 progeny were individually picked and screened for stable transmission of non-Unc, non-fluorescent progeny.

### Locomotion assays

*Crawling and radial locomotion assays* were performed using L4-staged hermaphrodite animals transferred onto 60 mm nematode growth medium (NGM) plates. After transfer onto plate, animals were given two minutes of recovery time before being recorded for two minutes using the Nikon AZ100 microscope with 1X PlanApo objectives and the Photometrics ES2 camera. The individual movement of worms in the videos were analyzed with NIS-Elements AR Analysis software. The coordinates of individual worms were tracked over a span of one minute to calculate radial distance and speed. The radial graph was plotted using a custom MatLab script to produce the radial dispersion graph.

*Swimming assays* were performed using L4 stage hermaphrodites transferred into a depression slide containing 1 ml M9 buffer as described (Miller *et al.* 1996; Opperman *et al.* 2015). After one minute of recovery time, the animals were recorded for one minute using Nikon AZ100 microscope with 1X PlanApo objectives and Photometrics ES2 camera. The number of times an animal thrashed (bending at mid body) was counted manually by examining the video recording.

All locomotion assays were performed with the experimenter blind to the genotype of each strain. To prevent outliers from skewing data, animals immobile for more than 10 seconds were removed from the data pool. All strains were grown on NGM media with OP50 bacteria at 20°C incubator.

### Isolation and mapping of *ksr-1(eq7)*

*Screen for genetic suppressors of the rab-3; sax-7 uncoordinated locomotion*. Synchronized L4-staged double mutant animals were incubated in 50 mM ethylmethylsulfonate for 4 hours at 20°C as described (Brenner 1974). Gravid F1 animals were placed at one distal end of 100 mm petri dishes with OP50 bacteria seeded on the opposite end. After 24 hours, we screened the F2 generation for animals that were able to traverse the plate to the bacterial lawn, indicating improved locomotion. From a small screen of 5,000 mutagenized haploid genomes, we isolated 4 suppressors. We characterized the strongest suppressor, *eq7,* mapping it to the linkage group X.

*Mapping of eq7*: *eq7* was further mapped using single nucleotide polymorphisms (SNPs) in the Hawaiian strain CB4856 as described (Davis *et al.* 2005). Co-segregation of SNPs with suppressed or non-suppressed animals was used to map *eq7* to a 250kb interval on LGX bounded by WBVar 00081116 and WBVar00054056. *ksr-1* was identified as a candidate gene within this interval; sequencing of the exons of the gene verified *eq7* as a nonsense mutation at codon 546. Additional confirmation was obtained via a complementation test with the *ksr-1(ok786)* deletion allele.

### Quantitative analysis of fluorescence microscopy

GABA and cholinergic synapses in motor neurons were examined using the respective SNB-1::GFP transgenes, *juIs1*, *wyIs92*, and *wdIs20* (Lickteig *et al.* 2001; Hallam *et al.* 2002; Klassen and Shen 2007). Young adult animals were mounted on 2% agarose pads and anesthetized using 10 mM levamisole in M9 buffer. Images were acquired with a Zeiss Axio Observer Z1 motorized microscope equipped with a C-ApoCHROMAT 63x/1.20 NA water immersion objective and a QuantEM512SC camera (Photometrics). The light source was an HXP-120 mercury halide lamp and the excitation and emission bands were 450-490 and 500-550 nm, respectively. Plane interval in Z acquisitions were 31 nm, for a voxel size of 254×254×31 nm. Acquisition software used was ZEN2 (Zeiss). To quantitate punctal fluorescence, images of the dorsal nerve cord were acquired by the posterior gonad bend. The images were then deconvolved with AutoQuantX software and sum slices projection was obtained using ImageJ. The average background intensity value was used to standardize the dorsal cord intensity. Punctal fluorescence values were scored over 100μm using the Find Maxima Function of ImageJ, n > 10 animals for each strain. All images were taken under the same magnification, binning, bit depth, and exposure time.

### Electrophysiology

Post-synaptic recordings were obtained from ventromedial body wall muscles anterior to the vulva from immobilized dissected worms, as previously described (Richmond and Jorgensen 1999). Briefly, spontaneous and electrically evoked synaptic responses were acquired from individual muscle cells, whole-cell voltage-clamped at a holding potential of −60mV in the presence of a 1mM Ca^2+^ extracellular solution, using a HEKA EPC-10 amplifier. Data were analyzed using Synaptosoft and Igor Pro.

### Electron Microscopy

Samples were prepared using the high-pressure freeze fixation and freeze substitution methods as previously described (Weimer 2006). ∼30 young adult hermaphrodites were placed in each specimen chamber containing *E. coli* and rapidly frozen to −180°C under high pressure (Leica HPM 100). Frozen specimens then underwent freeze substitution (Leica Reichert AFS) during which samples were held at −90°C for 107 hr in 0.1% tannic acid and 2% OsO4 in anhydrous acetone. The temperature was then raised at 5°C/hr to −20°C, kept at −20°C for 14 hr, and then raised by 10°C/hr to 20°C. After fixation, samples were infiltrated with 50% Epon/acetone for 4 hr, 90% Epon/acetone for 18 hr, and 100% Epon for 5 hr. Finally, samples were embedded in Epon and incubated for 48 hr at 65°C. Ultrathin (40 nm) serial sections were cut using a Leica Ultracut 6 and collected on formvar-covered, carbon-coated copper grids (EMS, FCF2010-Cu). Post-staining was performed using 2.5% aqueous uranyl acetate for 4 min, followed by Reynolds lead citrate for 2 min. Images were acquired using a JEOL JEM-1220 transmission electron microscope operating at 80 kV using a Gatan digital camera at a magnification of 100k (1.8587 pixels/nm). Images were collected from the ventral nerve cord region anterior to the vulva for all genotypes. Serial micrographs were manually annotated using NIH ImageJ/Fiji software to quantify the number of plasma membrane docked SVs, and distance of docked SVs from the dense projection (DP) for each synaptic profile. Specimens were encrypted to ensure unbiased analysis. Cholinergic synapses were identified on the basis of their typical morphology (White *et al.* 1986). A synapse was defined as a series of sections (profiles) containing a dense projection as well as two flanking sections on both sides without dense projections. SVs were identified as spherical, light grey structures with an average diameter of ∼30 nm. A docked synaptic vesicle was defined as a synaptic vesicle whose membrane was morphologically contacting the plasma membrane.

### Data and reagent availability

Strains and plasmids are available upon request. Supplemental material available at figshare.

## RESULTS

### *sax-7* interacts with SV cycle genes, resulting in a coiling behavior

We previously identified a genetic interaction between *sax-7* and *rab-3*, which encodes a monomeric G-protein that is associated with synaptic vesicles in its GTP-bound state (Opperman *et al.* 2015). *rab-3; sax-7* double mutant animals display synthetic uncoordinated (Unc), loopy locomotion with a tendency to coil that is not observed in either *rab-3* or *sax-7* single null animals (Fig 1A, S1 video). Although not paralyzed, *rab-3; sax-7* double mutants tend to meander locally, exhibiting apparent non-vectorial locomotion. As a result, *rab-3; sax-7* animals show severely limited radial displacement, in comparison to either *rab-3* or *sax-7* single mutant animals (Fig 1Bi, C).

**Fig 1.**
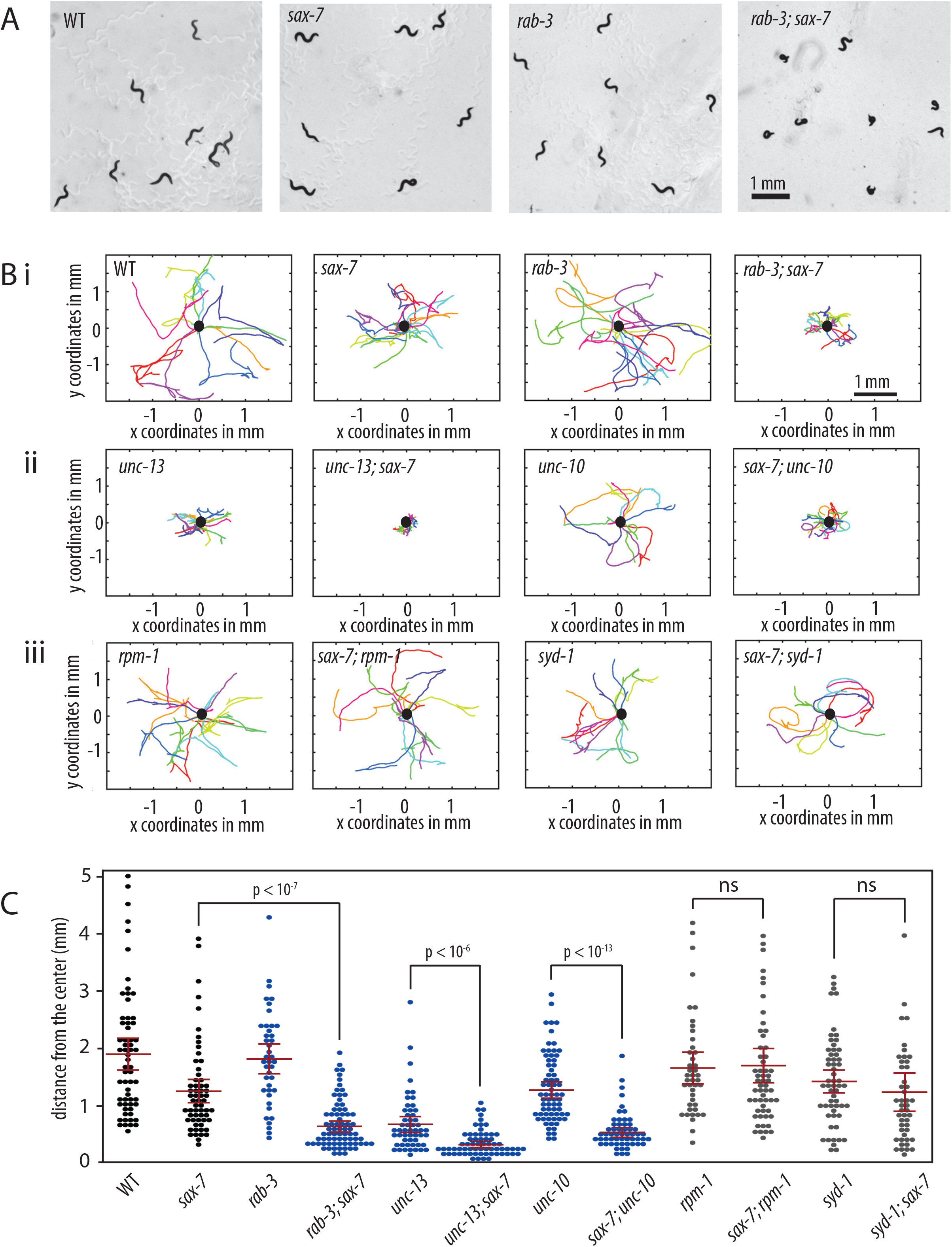
*sax-7* interacts with genes that function in the SV cycle but not in synaptogenesis. (A) *rab-3; sax-7* mutant animals exhibit synthetic Unc locomotion with a tendency to coil that is not observed in either *rab-3* or *sax-7* single mutant animals. (B) Graphs tracing the movements of 10 random animals per strain over a span of one minute. Each colored line represents the tracks of an individual animal with the point of origin marked as a black circle in the center (0,0 coordinate). These graphs reveal that *sax-7* interacts with SV cycle genes, (i) *rab-3* as well as (ii) *unc-13* and *unc-10,* resulting in a synergistic reduction in the ability to disperse. In contrast, *sax-7* does not interact with genes that function in synaptogenesis, (iii) *rpm-1* and *syd-1*. (C) The ability to disperse, quantified as the radial distance travelled by each animal over a span of one minute, is illustrated in a scatter plot where each point is a data point for a single animal; the mean and the 95% confidence intervals are marked in red. The blue plots are data points for mutant strains of SV cycle genes while grey plots show data points for mutant strains of synaptogenesis genes. n = 50 – 75, *p*-values are calculated using one-way ANOVA with Bonferroni’s *post hoc* test and compared between the double mutant and the single mutant exhibiting the stronger phenotype, n.s., not significant.

The RAB-3 protein promotes neurotransmission, acting in the synaptic vesicle (SV) cycle with roles that include the trafficking and docking of synaptic vesicles to the active zone at synapses (Nonet *et al.* 1997; Gracheva *et al.* 2008). The genetic interaction with *rab-3* suggests that *sax-7* may interact with additional genes that function in SV exocytosis. While the majority of SV cycle mutants exhibit severely Unc phenotypes that preclude enhancement, we were nonetheless able to test for an interaction with *unc-10,* which encodes for the RAB-3-interacting molecule (RIM) that recruits SVs to the active zone via RAB-3::GTP interactions (Koushika *et al.* 2001; Gracheva *et al.* 2008). We also confirmed a previously-identified interaction with *unc-13*, which functions in SV docking and priming at the active zone (Richmond *et al.* 1999; Weimer *et al.* 2006; Opperman *et al.* 2015). We observed synthetic coiling and poor dispersal behaviors in *sax-7* animals homozygous for either *unc-13* hypomorphic or *unc-10* null alleles (Fig 1Bii, C). These results are consistent with the notion that *sax-7* interacts with SV cycle genes. In contrast, we did not observe an interaction between *sax-7* and genes with roles in synaptogenesis, such as *rpm-1* and *syd-1* (Schaefer *et al.* 2000; Zhen *et al.* 2000; Hallam *et al.* 2002; Patel *et al.* 2006; Cherra and Jin 2015). Loss of either *rpm-1* or *syd-1* in *sax-7* null animals does not produce the coiling and poor dispersal behavior exhibited by *rab-3; sax-7* animals (Fig 1Biii, C). These results indicate that the *sax-7* interaction with SV cycle genes is specific.

The abnormal locomotory behaviors in double mutants of *sax-7* and SV cycle genes suggest underlying neuronal dysfunction. To test this notion, we measured the ability of *rab-3; sax-7* animals to swim in liquid and crawl on agar medium. The ability for animals to swim was quantified as the number of times an animal thrashes in liquid per minute. While wild-type animals have an average swim rate of 163 thrashes/minute, the mean swim rate of *rab-3; sax-7* animals is 33 thrashes/minute, which is only 20% of the wild-type rate (Fig 2A). By comparison, the average swim rates of *sax-7* and *rab-3* single mutants are 53% and 85% that of wild-type, respectively. *rab-3; sax-7* double mutants also show a synergistic decrease in crawl rates (Fig 2B). At 35 μm/second, *rab-3; sax-7* animals crawl at an average rate that is 44% of the wild-type rate (80.5 μm/second) whereas the average crawl rates of *sax-7* and *rab-3* single mutants are 82% and 116% of the wild-type rate, respectively. Synergistic reductions in swim and crawl rates are also observed in *sax-7* double mutant animals of *unc-13, unc-10*, and *snb-1*, which encodes synaptobrevin (Nonet *et al.* 1998). In contrast, *sax-7* double mutants with synaptogenesis genes, *rpm-1* and *syd-1,* exhibit modest, if any, synergism (Fig 2). These results are not only consistent with neuronal dysfunction contributing to the synthetic Unc and locomotion behaviors, but also underscore the specificity of the *sax-7* genetic interaction with SV cycle genes.

**Fig 2.**
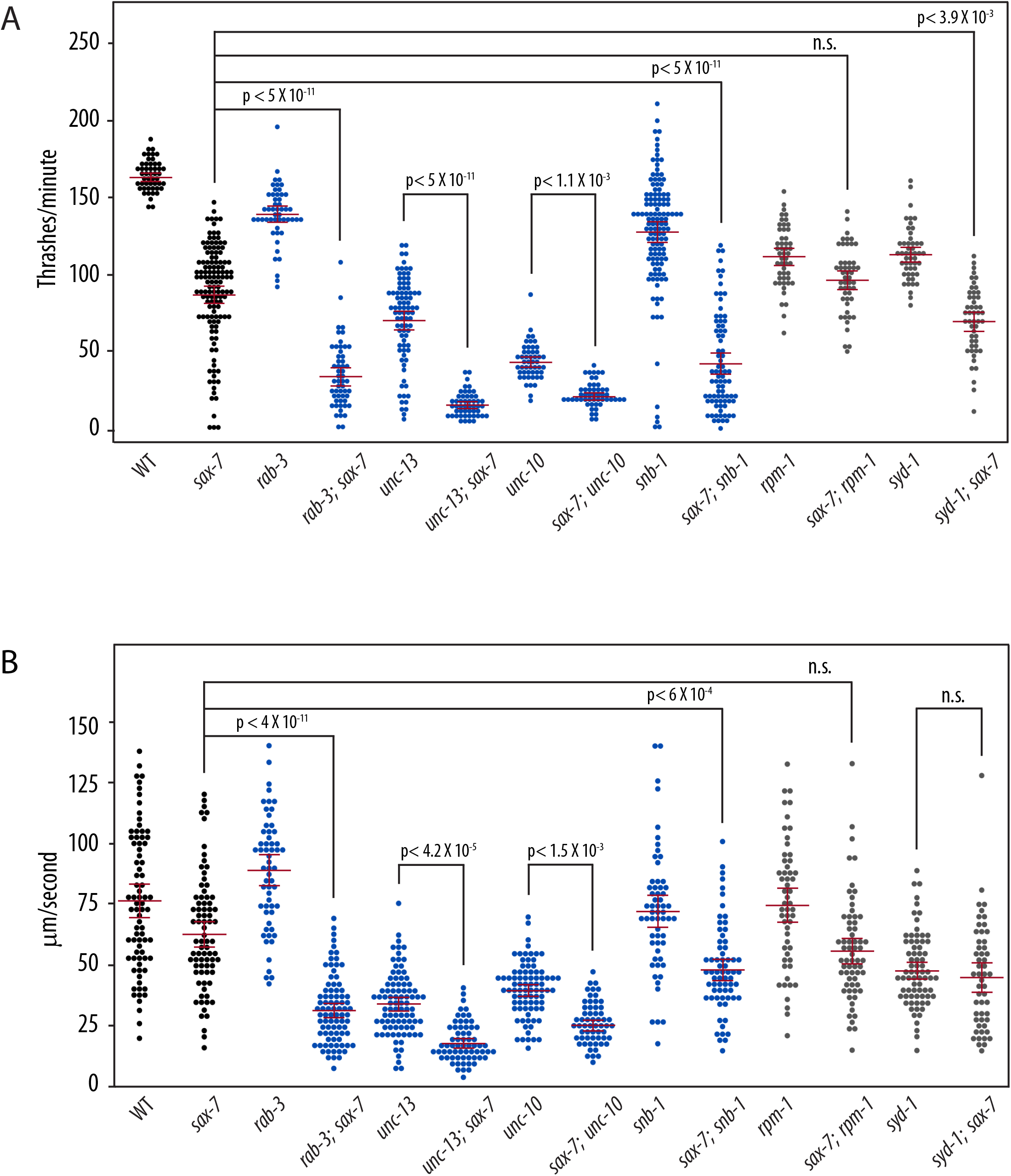
*sax-7* double mutants with SV cycle genes display synergistic neuronal dysfunction. (A) sax-7 interacts with SV cycle but not synaptogenesis genes, showing synergistic reduction in swim rates (number of thrashes per minute). (B) sax-7 interacts with SV cycle but not synaptogenesis genes, resulting in a synergistic reduction in crawl rates (μm per second). The blue plots are data points for mutant strains of SV cycle genes while grey plots show data points for mutant strains of synaptogenesis genes. The mean and the 95% confidence intervals are marked in red. n = 50-75, *p*-values are calculated using one-way ANOVA with Bonferroni’s *post hoc* test and compared between the double mutant and the single mutant exhibiting the stronger phenotype, n.s., not significant.

### *Sax-7* is required in the nervous system to promote coordinated locomotion

*sax-7* is expressed both in the nervous system and non-neuronal tissues, including body-wall muscles and the hypodermis (Chen *et al.* 2001). Based on the locomotory deficiencies observed in *sax-7* double mutants with SV cycle genes, we predicted that *sax-7* expression in the nervous system would suppress *rab-3; sax-7* phenotypes. However, non-neuronal *sax-7* expression has been shown previously to impact the maintenance of neural architecture, dendrite morphogenesis, and axon branching (Wang *et al.* 2005; Dong *et al.* 2013; Salzberg *et al.* 2013; Diaz-Balzac *et al.* 2016; Zhu *et al.* 2017). To determine where *sax-7* is required for coordinated locomotion, we first established a conditional *sax-7* knock-in system using the Crispr-Cas9-engineered *sax-7*(*eq22*) allele. *eq22* is an in-frame DNA insertion in the penultimate exon of the *sax-7* gene consisting of coding sequence for mCherry that is interrupted by a gene-disrupting cassette comprising *Pmyo-3::GFP::unc-54 3’UTR* and *Prps-27::neoR::unc-54 3’UTR* sequences (Fig 3A, Norris *et al.* 2015). As such, we expect disrupted *sax-7* function in *sax-7(eq22)* animals, which can be identified as neomycin-resistant animals that express GFP in pharyngeal muscles. Indeed, *sax-7(eq22)* animals resemble *sax-7(eq1)* null animals, showing comparable reduced swim rates of 89.6 thrashes/minute (Fig 3B). Importantly, *eq22* also genetically interacts with *rab-3*, producing synthetic coiling and Unc locomotion as well as a synergistically low swim rate similar to that observed in *rab-3; sax-7(eq1)* animals (Fig 3B). These results are consistent with *eq22* disrupting *sax-7* function and further demonstrate that the *sax-7* genetic interaction we observed with *rab-3* is not allele-specific.

**Fig 3.**
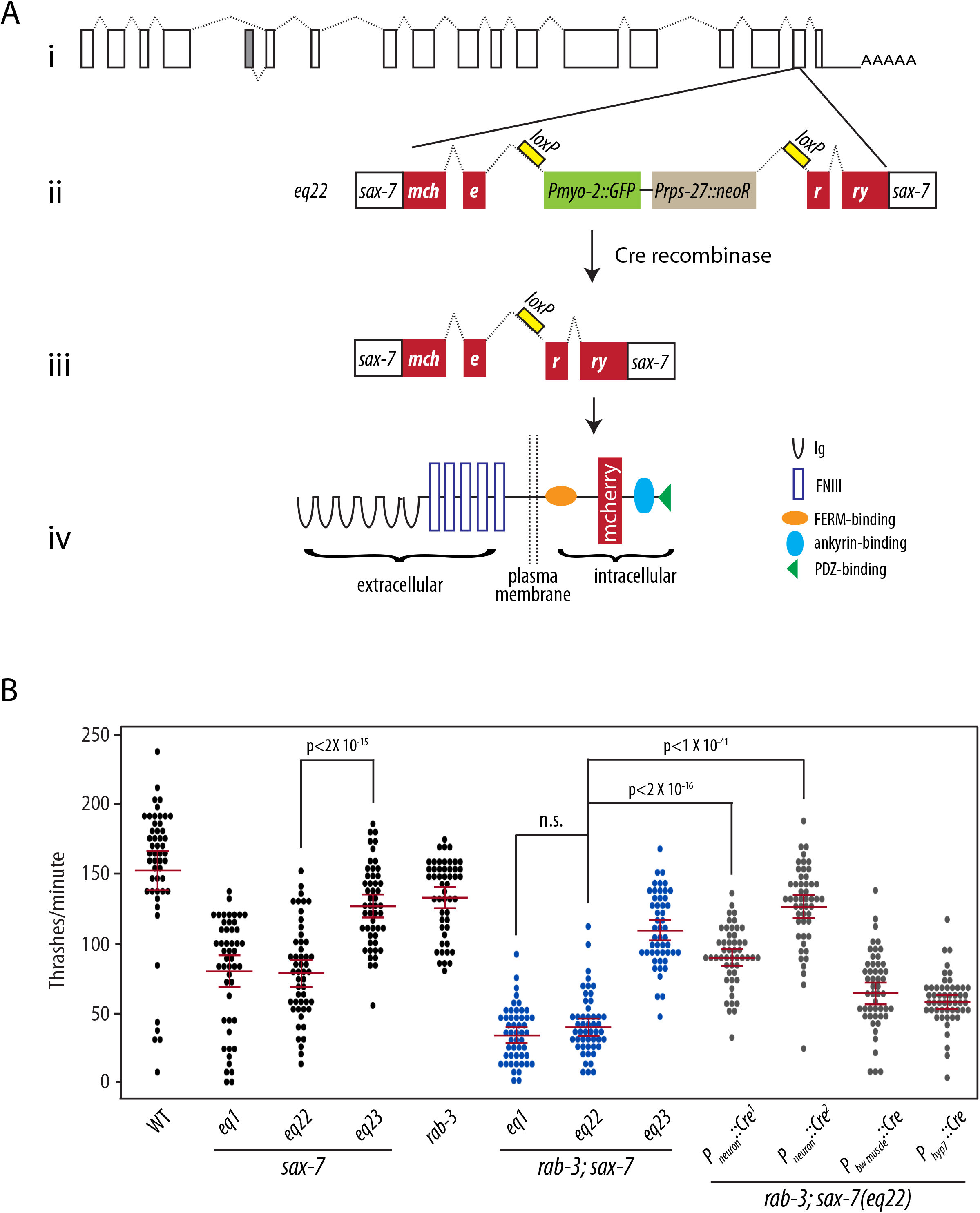
Neuronal expression of *sax-7* rescues *rab-3; sax-7* uncoordinated locomotion and neuronal dysfunction. (A) A schematic of the conditional *sax-7* knock-in allele, *eq22*. (i) Gene structure of the *sax-7* locus, with boxes representing exons and dotted lines representing introns; the grey box represents an exon used in an alternatively-spliced isoform of *sax-7* (Chen *et al.* 2001)*. eq22* is an in-frame insertion into the penultimate *sax-7* exon (ii). The insertion contains mCherry sequence and a gene-disrupting cassette (P*myo-2*::GFP and P*rps-27*::neoR) that interrupts the mCherry reading frame. This gene-disrupting cassette, which is flanked by loxP sites located in intronic sequences, can be excised out with tissue-specific Cre-recombinase expression (iii) to restore *sax-7* expression as (iv) a full-length SAX-7::mCherry fusion protein. Cre-recombinase expression in the germline was used to generate the *eq23* allele. (B) Quantitation of thrash rates reveal that *eq22* resembles the *eq1* null allele and similarly interacts with *rab-3* for a synergistically reduced swim rate; in contrast, *eq23* does not interact with *rab-3*. Conditional knock-in of *sax-7* in neurons with pan neuronally-expressed Cre-recombinase dramatically rescues the low *rab-3; sax-7*(*eq22*) thrash rate; in contrast, body-wall muscle or hypodermal knock-in results in modest rescue. Pan neuronally-expressed Cre-recombinase, P_neuron_::Cre^1^ and P_neuron_::Cre^2^, was produced using the *eqIs6* and *tmIs778* transgenes, respectively. The red lines show the mean and the 95% confidence interval. n = 50-75, *p*-values are shown, n.s., not significant, one-way ANOVA with Bonferroni’s *post hoc* test.

In addition to the gene-disrupting cassette, the *eq22* DNA insertion also contains *loxP* sites flanking the gene-disrupting cassette (Fig 3Aii). The *loxP* sites provide the ability to direct the excision of the gene-disrupting cassette with tissue-specific Cre-recombinase expression (Fig 3Aiii) for a targeted restoration of SAX-7 as a mCherry fusion protein (Fig 3Aiv). Using germline Cre-recombinase expression, we successfully excised the gene-disrupting cassette in the *sax-7(eq22)* germline to generate the *eq23* allele. *sax-7(eq23)* animals are not neomycin-resistant and do not express GFP in the pharynx. Instead, they show SAX-7::mCherry expression and localization (Fig S2A) similar to that of endogenous SAX-7 (Chen *et al.* 2001). Importantly, *sax-7(eq22)* phenotypes are suppressed in *sax-7(eq23)* animals (Fig 3B). For example, *sax-7(eq23)* animals display an increased average swim rate of 144.75 thrashes/minute as compared to the mean *sax-7(eq22)* swim rate of 89.6 thrashes/minute. As expected, *rab-3; sax-7(eq23)* do not display Unc locomotion or the synergistically-low swim rate that are exhibited in both *rab-3; sax-7(eq1)* and *rab-3; sax-7(eq22)* animals. These results indicate that the SAX-7::mCherry protein expressed in *sax-7(eq23)* animals is functional.

To assess whether SAX-7 is required in the nervous system for proper locomotion in a *rab-3* background, we introduced the *eqIs4* transgene, which directs neuronal Cre-recombinase expression with the *rab-3* promoter, into *rab-3; sax-7(eq22)* animals. In *rab-3; sax-7(eq22); eqIs4* animals, neuronal *sax-7* expression is recovered with mCherry fluorescence present only in the nervous system (Fig S2B). Importantly, *eqIs4* suppresses the *rab-3; sax-7(eq22)* Unc behavior and low swim rates, as does another neuronal-expressing Cre transgene, *tmIs778,* which directs Cre-recombinase expression with the *rgef-1* promoter (Fig 3B (Kage-Nakadai *et al.* 2014)). The *rab-3; sax-7(eq22)* mean swim rate of 39.5 thrashes/minute is restored to a mean rate of 89.7 thrashes/minute by *eqIs4* and 126 thrashes/minute by *tmIs778*. In contrast, the low *rab-3; sax-7(eq22)* thrash rate is not rescued with the recovery of SAX-7 expression in either body-wall muscles (Fig S2C) or hypodermis (not shown) using the *tmIs1087* and *tmIs1028* transgenes, respectively (Fig 3 (Kage-Nakadai *et al.* 2014)). These results demonstrate a requirement for SAX-7 in the nervous system for coordinated locomotion in a *rab-3* null background.

### A role for Erk signaling in locomotory behavior and function

We performed a forward genetic screen for suppressors of the *rab-3; sax-7* Unc and coiling behavior as an approach to determine the basis of the abnormal *rab-3; sax-7* locomotion and neuronal function. We isolated a strong suppressor, *eq7*, a C-to-T transition that converts codon 546 of the *ksr-1* gene into a nonsense mutation (see materials and methods). *rab-3; sax-7; ksr-1(eq7)* triple animals display similar locomotory and dispersal behaviors as wild-type animals (Fig 4, S3A video). *ksr-1(eq7)* also rescues the abnormal *rab-3; sax-7* mean swim rates from 33 thrashes/minute to 121 thrashes/minute as well as crawl rates from 35μm/second and 67μm/second (Fig 5). The *ksr-1* deletion allele, *ok786* (Moerman and Barstead 2008; Consortium 2012), similarly suppresses *rab-3; sax-7* phenotypes (Fig 4, 5). These results are consistent with the loss of *ksr-1* function suppressing *rab-3; sax-7* abnormal locomotion and neuronal defects. *ksr-1* single mutants do not have apparent locomotion deficiencies, displaying thrash and crawl rates that are similar or modestly reduced as compared to wild-type animals.

**Fig 4.**
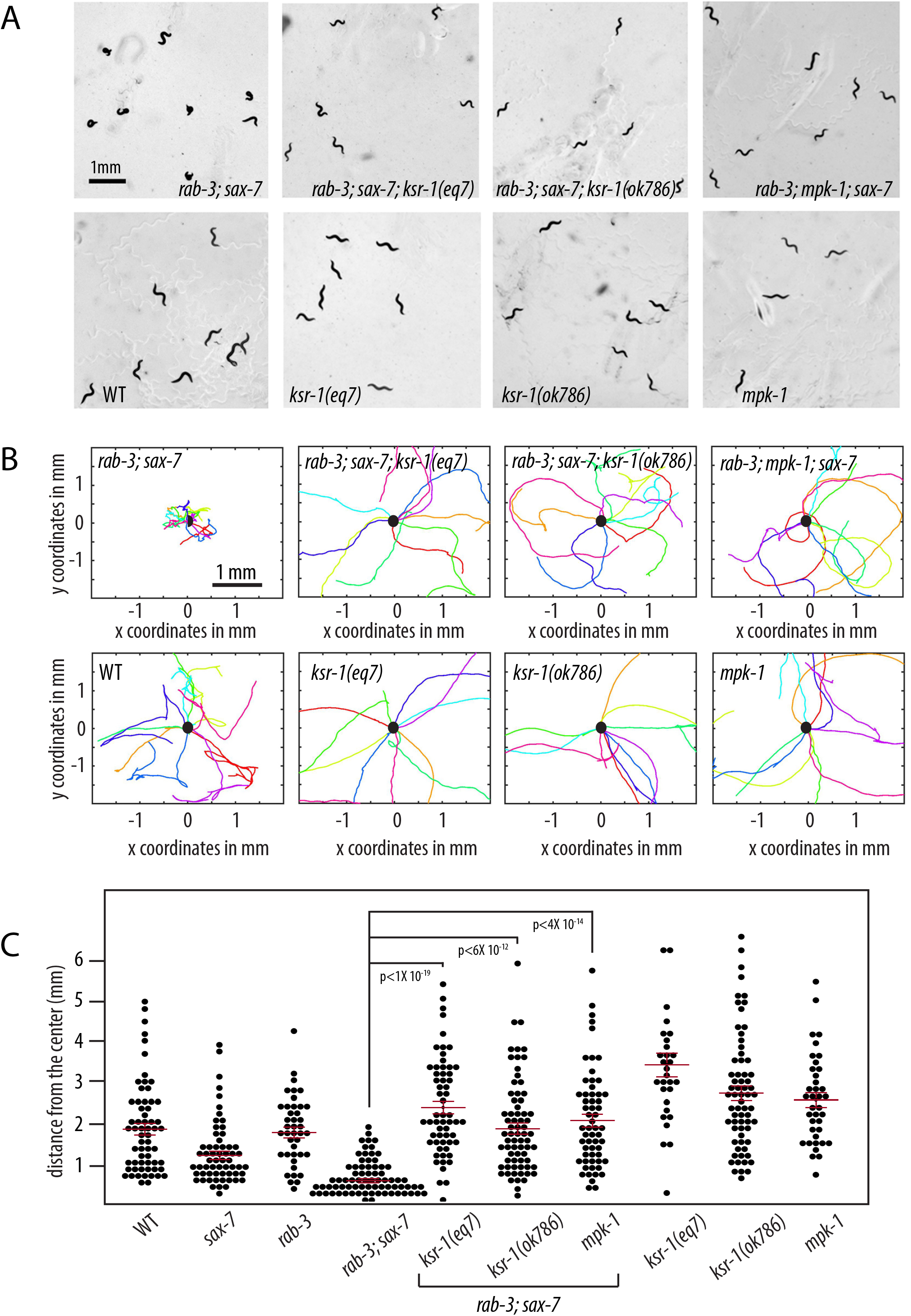
Reducing Mitogen Activated Protein Kinase (MAPK) signaling suppresses *rab-3; sax-7* uncoordinated locomotion. (A) Null alleles of *ksr-1* or *mpk-1* suppress *rab-3; sax-7* uncoordinated locomotion; yet *ksr-1* or *mpk-1* null animals themselves are not Unc. (B) Graphs tracing the movements of 10 random animals per strain over a span of one minute. Each colored line represents the tracks of an individual animal with the point of origin marked as a black circle in the center (0,0 coordinate). These graphs reveal that loss of *ksr-1* or *mpk-1* effectively suppress *rab-3; sax-7* reduced ability to disperse. (C) The ability to disperse, quantified as the radial distance travelled by each animal over a span of one minute, is illustrated in a scatter plot where each point is a data point for a single animal; the mean with the 95% confidence interval are marked in red. n = 50 – 75, *p*-values are shown, n.s., not significant, one-way ANOVA with Bonferroni’s *post hoc* test.

**Fig 5.**
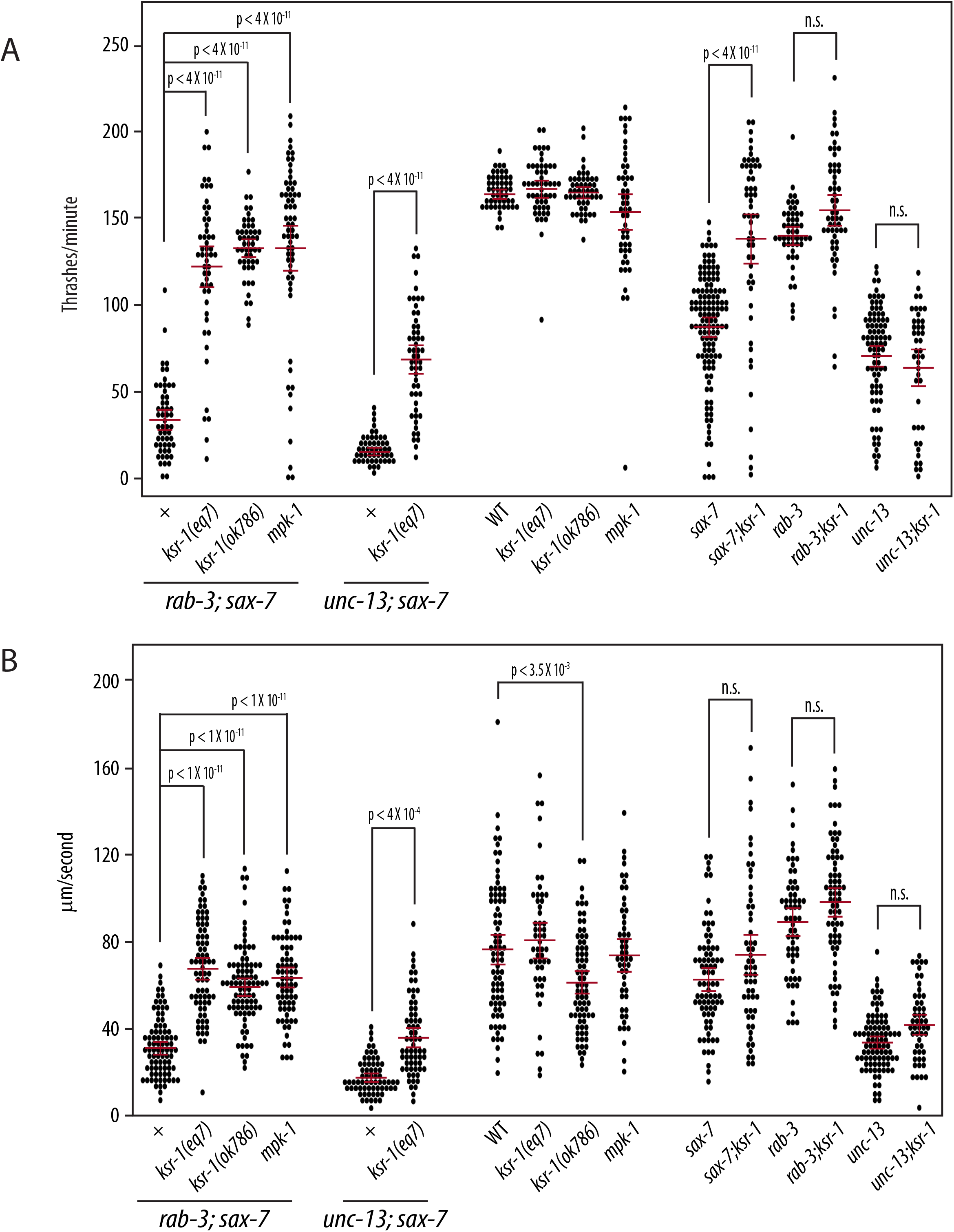
Reduced MAPK signaling suppresses *sax-7-*related neuronal dysfunction. Null alleles of *ksr-1* and *mpk-1* can suppress both *rab-3; sax-7* and *unc-13; sax-7* synergistically reduced (A) swim and (B) crawl rates; yet *ksr-1* or *mpk-1* null animals have similar swim and crawl rates as wild-type animals. Loss of *ksr-1* can also suppress the reduced swim rate exhibited by *sax-7* null animals, but not *unc-13* mutants, suggesting that the suppression of *rab-3; sax-7* and *unc-13; sax-7* neuronal dysfunction is due to *ksr-1* impinging on the *sax-7* pathway. The mean with the 95% confidence interval are marked in red. n = 50-75. n.s., not significant, one-way ANOVA with Bonferroni’s *post hoc* test.

In addition to *rab-3; sax-7* phenotypes, *ksr-1(eq7)* also suppresses *unc-13; sax-7* defects, including the low swim and crawl rates (Fig 5). This suppression in the double mutants raises the possibility that *ksr-1* may act on the *sax-7* pathway, the SV cycle, or function in a third as-yet-unidentified pathway promoting coordinated locomotion and neuronal function. Interestingly, the rescued *rab-3; sax-7; ksr-1* and *unc-13; sax-7; ksr-1* swim and crawl rates are more similar to the *rab-3* and *unc-13* rates, respectively (Fig 5), suggesting that the loss of *ksr-1* may be impinging on the *sax-7* pathway rather than the SV cycle. If this hypothesis is correct, loss of *ksr-1* should suppress *sax-7* phenotypes but not affect *rab-3* or *unc-13* phenotypes. This hypothesis is best tested by examining the swim rates rather than crawl rates since the *sax-7* phenotype is more robust in liquid than on solid media (Fig 2, 5). In liquid, the *sax-7* swim rate of 86 thrashes/minute is 51% of wild-type while the *sax-7; ksr-1* swim rate of 154 thrashes/minute is 94% of the wild-type rate, showing almost complete suppression by loss of KSR-1. By comparison, the suppression on *unc-13* swim rate by loss of KSR-1 is negligible (Fig 5A). In addition, the relatively normal swim rate of *rab-3* single mutants is not impacted by the loss of KSR-1 (Fig 5A). This genetic interaction observed between *ksr-1* and *sax-7* but not *unc-13* or *rab-3* is consistent with the notion that KSR-1 likely affects SAX-7 function.

KSR-1 is a molecule that interacts with core components of the Mitogen-Activated Protein Kinase (MAPK) cascade to facilitate the activation of the MAPK signaling pathway (Sundaram and Han 1995; Sundaram 2013). As both putative loss-of-function *ksr-1* alleles suppressed *rab-3; sax-7* phenotypes, we predicted that reducing the function of the MAPK, Erk, would similarly suppress the *rab-3; sax-7* phenotypes. To test this prediction, we crossed a null allele of Erk, encoded by the *mpk-1* gene (Lackner *et al.* 1994), into *rab-3; sax-7* animals. Consistent with our hypothesis, *rab-3; mpk-1; sax-7* triple mutant animals display locomotory behavior and ability to swim that are similar to that of wild-type animals (Fig 4, 5, and S3B video), revealing a role for MAPK signaling in regulating locomotion and neuronal function. Interestingly, both *ksr-1* and *mpk-1* single mutant animals do not display apparent locomotory deficiencies, suggesting that the role the MAPK signaling pathway plays in coordinating locomotion is likely modulatory.

### SAX-7 and KSR-1 do not affect SV release at neuromuscular junctions

A delayed response to the paralytic effects of the cholinesterase inhibitor, aldicarb, is often observed in animals with impaired SV exocytosis due to a slower build-up of acetylcholine in the synaptic clefts of neuromuscular junctions, as compared to animals with wild-type SV exocytosis (Miller *et al.* 1996). Consistent with the role of *rab-3* in the trafficking and recruitment of SVs to the active zone, *rab-3* mutant animals are mildly **r**esistant to **i**nhibitors of **c**holinesterase (RIC) (Nonet *et al.* 1997; Gracheva *et al.* 2008). Although *sax-7* animals are not RIC, loss of SAX-7 synergistically enhances the RIC phenotype in *rab-3* animals (Opperman *et al.* 2015). The *rab-3; sax-7* RIC phenotype, taken together with the accompanying Unc behavior and reduced swim and crawl rates, suggests that *rab-3; sax-7* animals may have synergistic SV exocytosis defects.

To resolve potential synergistic SV release abnormalities underlying the *rab-3; sax-7* phenotypes, we performed electrophysiological analyses of neuromuscular synapses to assess functional consequences in *rab-3; sax-7* double as compared to *rab-3, sax-7,* and *ksr-1* single mutant animals as well as the *rab-3; sax-7; ksr-1* triple mutant. Whole-cell voltage-clamped body recordings from medial ventral body wall muscles were directly assayed before and after electrical stimulation of the ventral nerve cord. First, the average endogenous miniature event amplitude observed in each genotype is similar to that of wild-type, suggesting that postsynaptic muscle receptor function does not account for the phenotypes observed in *rab-3; sax-7* animals. This finding is consistent with the inability of SAX-7 expression in muscle to rescue *rab-3; sax-7* phenotypes (Fig 3B). Second, we examined the evoked responses in each strain. As expected, *rab-3* null animals exhibit reduced evoked responses, consistent with previous studies (Nonet *et al.* 1997; Gracheva *et al.* 2008), while *sax-7* and *ksr-1* animals showed wild-type evoked responses. Surprisingly and contrary to our prediction, *rab-3; sax-7* double mutants evoked responses that were not significantly lower than that of *rab-3* mutants (Fig 6). In fact, there was no significant difference among the evoked responses of *rab-3* single*, rab-3; sax-7* double, and *rab-3; sax-7; ksr-1* triple mutant animals.

**Fig 6.**
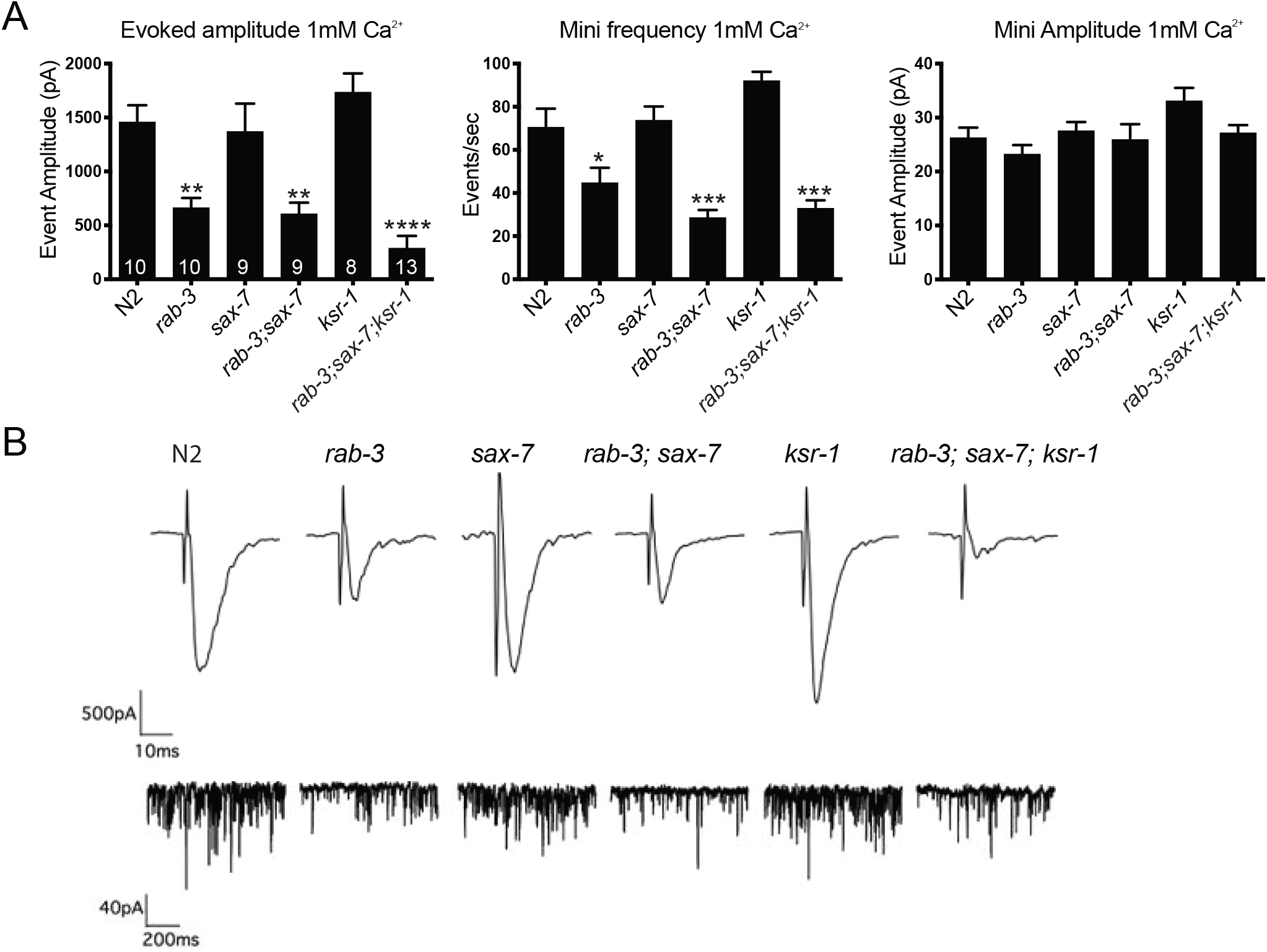
SAX-7 and KSR-1 do not affect neurotransmitter release in ventral cord cholinergic motor neurons. (A) Whole-cell voltage-clamped body recordings from medial ventral body wall muscles (held at −60mV in 1mM Ca^2+^) of dissected worms. The graphs show the average evoked amplitude, mini frequency, and mini amplitude at the NMJ of each strain. The number of animals analyzed per genotype is shown within each bar. Data shown as mean ± SEM. (B) Representative evoked traces (top panel) and miniature traces (bottom panel) for genotypes labeled on the top. *P*-values are calculated using one-way ANOVA with Tukey’s multiple comparison test comparing the different strains to wild-type; \**P* < 0.05, \*\**P* < 0.01, \*\*\**P* < 0.0001

We also examined *rab-3; sax-7* synapses at the ultrastructural level using high pressure freeze electron microscopy (HPF-EM). As a control, we first compared *rab-3* cholinergic presynaptic terminals with wild-type. Consistent with published studies, we observed reduced numbers of docked SVs within 100nm of the presynaptic density in *rab-3* synapses (Gracheva *et al.* 2008). Interestingly, *rab-3; sax-7* synapses displayed similar SV numbers docked within 100nm of the presynaptic density as *rab-3* synapses, while *sax-*7 synapses showed wild-type SV numbers (Fig 7). Moreover, *rab-3; sax-7; ksr-1* triple mutant animals showed a similar docking deficit as *rab-3* single and *rab-3; sax-7* double mutant animals. Surprisingly, *ksr-1* synapses also presented a deficit in vesicle docking that is not consistent with the suppression of *rab-3; sax-7* neuronal phenotypes. These results are consistent with our findings obtained by examining the synapses using light microscopy. Using GFP-tagged synaptobrevin (SNB-1::GFP) as a fluorescence marker for synaptic vesicles, we determined there was no apparent difference in SNB-1::GFP localization, number, or punctal intensity in *rab-3; sax-7* GABA and cholinergic synapses as compared to wild-type (Fig S4). In contrast, the punctal fluorescence intensity of SNB-1::GFP is increased in *unc-18* synapses as expected when SV release is impaired; *unc-18* encodes for a syntaxin-binding protein necessary for SV exocytosis (Richmond 2005; Ch’ng *et al.* 2008). Taken together, these data indicate that altered presynaptic function at the ventral cord cholinergic neuromuscular junctions is unlikely a contributing cause for the *rab-3; sax-7* locomotory behaviors and neuronal dysfunction, and cannot explain the ability of *ksr-1* mutants to suppress these defects. This finding suggests that *ksr-1* suppression of the *rab-3; sax-7* phenotype occurs elsewhere in the nervous system.

**Fig 7.**
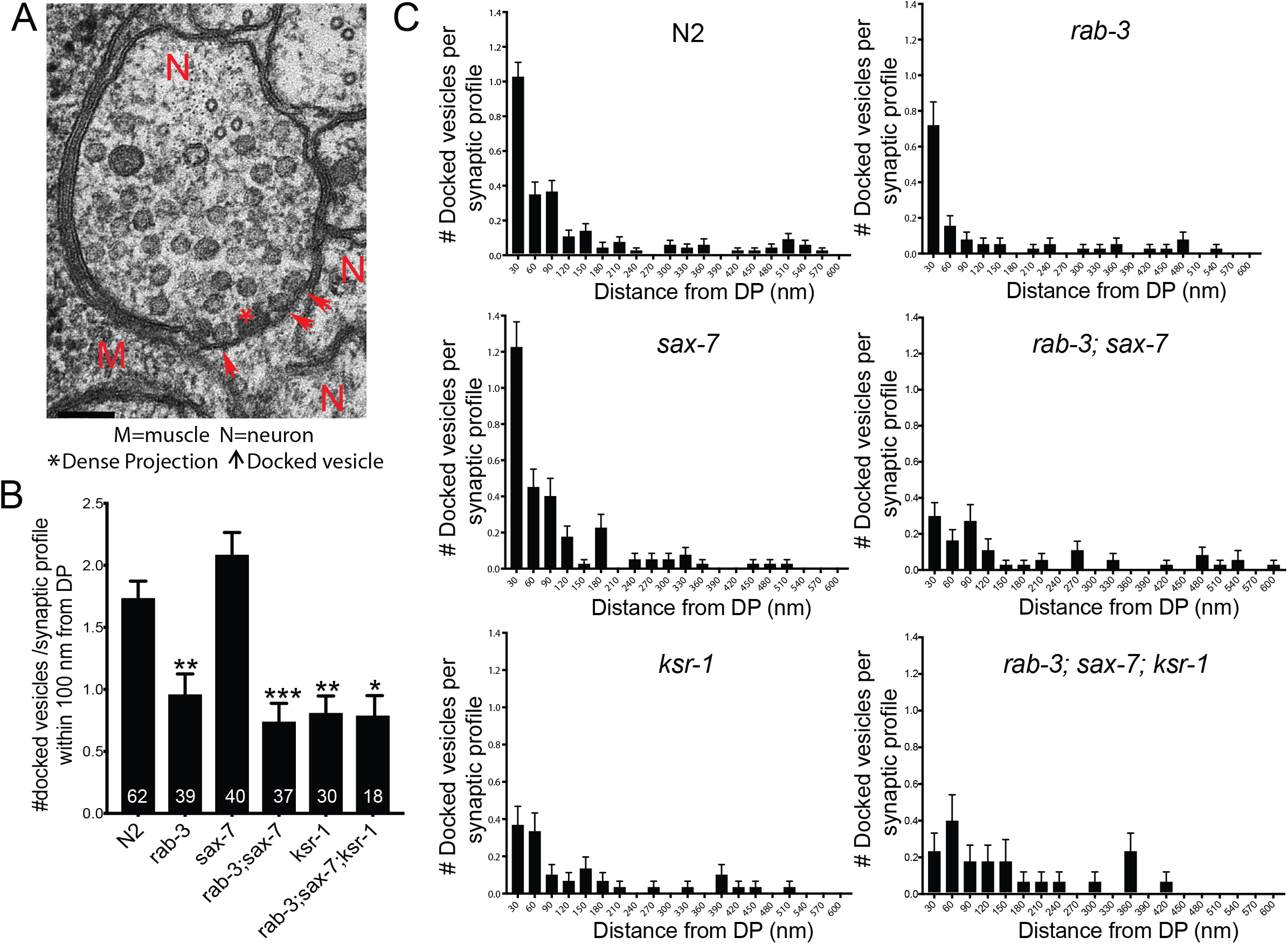
SAX-7 and KSR-1 do not affect number or distribution of docked SVs in ventral cord cholinergic motor neurons. (A) A representative electron micrograph of a cholinergic synaptic profile. (B) Average number of docked SVs per synaptic profile within 100 nm distance from dense projection (DP) or presynaptic density. Number of synaptic profiles analyzed per genotype is mentioned within the bars. (C) Distribution of docked SVs with respect to the dense projection. Data shown as mean ± SEM, *p*-values are calculated using one-way ANOVA with Tukey’s multiple comparison test comparing the different strains to wild-type; \**P* < 0.05, \*\**P* < 0.01, \*\*\**P* < 0.0001

### *ksr-1* functions in a subset of cholinergic neurons in the head for coordinated locomotion

A recent study uncovered a novel neuronal role for *ksr-1* (Coleman *et al.* 2018). Specifically, loss-of-function *ksr-1* alleles were isolated in a genetic screen for suppressors of the hyperactive and loopy locomotion exhibited by activated Gq/EGL-30 animals. Moreover, *ksr-1* expression restricted to a subset of cholinergic neurons in the head of *egl-30; ksr-1* animals was sufficient to reverse this suppression; in contrast, *ksr-1* expression in ventral nerve cord cholinergic neurons in the body did not (Coleman *et al.* 2018).

Because of overlapping loopy locomotion shared by both *egl-30* and *rab-3; sax-7* animals, we hypothesized that *ksr-1* might similarly act in cholinergic neurons to regulate *rab-3; sax-7* locomotion. If so, *ksr-1* expression in cholinergic neurons should reverse the suppression observed in *rab-3; sax-7; ksr-1* animals. Indeed, using the *eqSi3* transgene to drive single-copy transgenic KSR-1 expression under the neuronal *unc-17* acetylcholine transporter promoter, *rab-3; sax-7; eqSi3; ksr-1* animals strongly resemble *rab-3; sax-7* animals, exhibiting reduced swim rates as well as Unc, loopy locomotion with a tendency to coil (Fig 8A). Moreover, using the *eqSi1* single-copy transgene to restrict transgenic KSR-1 expression to cholinergic neurons in the head with the partial *unc-17H* promoter was also sufficient to reverse the suppression observed in *rab-3; sax-7; ksr-1* animals. Indeed, *rab-3; sax-7; eqSi1; ksr-1* animals are Unc, displaying coiling locomotion and reduced swim rates (Fig 8A). In contrast, using the *eqSi2* single-copy transgene to limit *ksr-1* expression to cholinergic neurons in the body with the *unc-17B* promoter *was* not sufficient to reverse the suppression in *rab-3; sax-7; ksr-1* animals. Importantly, the same single-copy transgenes did not impact locomotion or swim rate in wild-type animals (Fig 8A). These results indicate that the *ksr-1* expression in the head cholinergic neurons is sufficient to promote both coordinated locomotion and neuronal function, and likely account for the absence of neuronal dysfunction or abnormality in ventral nerve cord synapses assayed in our electrophysiological and ultrastructural analyses.

**Fig 8.**
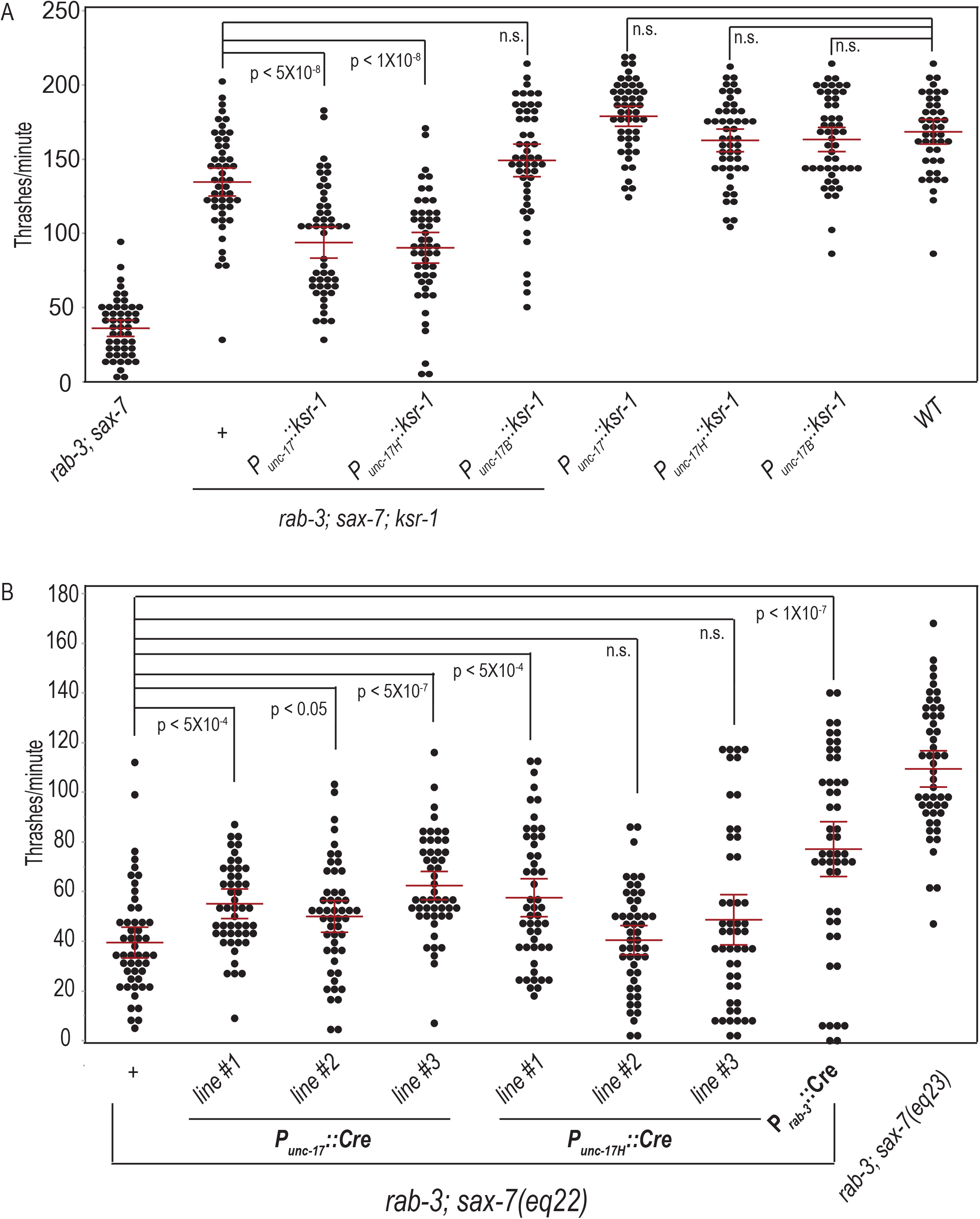
*ksr-1* functions in a subset of cholinergic neurons in the head to modulate *rab-3; sax-7* neuronal function. A) The loss of *ksr-1* suppresses *rab-3; sax-7* reduced swim rate. This suppression is reversed when *ksr-1* is expressed in *rab-3; sax-7; ksr-1* cholinergic neurons using *unc-17* acetylcholine transporter promoter. This reversal is similarly observed when *ksr-1* expression is restricted to cholinergic neurons in the head using the partial *unc-17H* promoter; in contrast, no reversal is observed when *ksr-1* expression is confined to body cholinergic motor neurons using the partial *unc-17B* promoter. Wild-type animals carrying each transgene exhibit normal swim rates and show no apparent locomotion phenotype. B) To determine whether *sax-7* functions in the same neurons as *ksr-1*, we used the conditional *sax-7*(*eq22*) knock-in allele and P*_unc-17_*::Cre-recombinase to knock in *sax-7* expression in cholinergic neurons. *sax-7* expression in all or only head cholinergic neurons shows modest or poor suppression of *rab-3; sax-7* decreased swim rates. The mean with the 95% confidence interval are marked in red. n = 50-75. n.s., not significant, one-way ANOVA with Bonferroni’s *post hoc* test.

We next assessed whether *sax-7* functions in the same set of neurons as *ksr-1* in coordinating locomotion. Using the same *unc-17* promoter, we directed Cre-recombinase in *rab-3; sax-7(eq22)* cholinergic neurons animals from multicopy transgenes on extrachromosomal arrays to restore *sax-7* expression. In three independent transgenic lines, we did not observe suppression of the Unc locomotion; however, there was a modest but significant suppression of the mean swim rate that ranged from 50-62 thrashes/minute as compared to the mean *rab-3; sax-7(eq22)* rate of 39.5 thrashes/minute (Fig 8B). Similarly, restoring *sax-7* expression in head cholinergic neurons resulted in little, if any, suppression of the *rab-3; sax-7*(*eq22*) swim rate. By comparison, extrachromosomal arrays (Fig 8B) as well as single-copy transgenes (Fig 3B) used to drive Cre-recombinase pan-neuronally resulted in a stronger suppression of the *rab-3; sax-*7(*eq22*) swim rate. These results indicate that *sax-7* expression in cholinergic neurons alone is not sufficient to rescue *rab-3; sax-7* phenotypes, suggesting *sax-7* function is also required in additional non-cholinergic neurons.

## Discussion

We previously uncovered a role for *sax-7* in promoting coordinated locomotion, first revealed in sensitized genetic *rab-3* and *unc-13* mutant backgrounds, in which *sax-7* mutants produced a synergistic decrease in locomotion. Adult-onset, transient *sax-7* expression could suppress the synergistic defects, consistent with a post-developmental role for the SAX-7/L1CAM protein. On the other hand, this transient *sax-7* expression did not suppress the well-established *sax-7* defect in maintaining neuronal and axonal positioning, suggesting a novel role for *sax-7* in promoting coordinated locomotion and neuronal function (Opperman *et al.* 2015). In this study, we examined these genetic interactions further and found them to be specific to genes that encode key players in the SV cycle necessary for neurotransmission. In contrast, genes that function in synapse formation do not interact with *sax-7* (Fig 1, 2). In addition to abnormal locomotion with a tendency to coil, *sax-7* animals in sensitized backgrounds of SV cycle mutants showed synergistic neuronal dysfunction manifested as decreased crawl and swim rates (Fig 1, 2). This *sax-7* role in promoting coordinated locomotion and appropriate swim and crawl rates is dependent on *sax-7* expression in the nervous system (Fig 3). We further demonstrate that this SAX-7 function in promoting coordinated locomotory behaviors is regulated by KSR-1 and MPK-1 (Fig. 4).

In addition to the specific interaction with SV cycle genes, we previously showed *rab-3; sax-7* animals also displayed strong resistance to the cholinesterase inhibitor, aldicarb, reflecting impaired cholinergic signaling (Opperman *et al.* 2015). Taken together, these findings suggested impaired synaptic activity as underlying these phenotypes and a potential role for *sax-7* in synaptic modulation. However, the electrophysiological recordings and ultrastructural analyses at neuromuscular junctions that were performed in this study did not reveal presynaptic abnormalities that could account for the observed behavioral phenotypes. Specifically, the cholinergic release defects associated with *rab-3* mutants were not exacerbated by loss of *sax-7*, nor was the release defect of *rab-3; sax-7* double mutants suppressed in the absence of *ksr-1*. Taken together, these findings suggest a potential role for SAX-7 and KSR-1 in synaptic modulation, but this does not appear to be at the level of the ventral cord neuromuscular junctions.

Our study also identified the MAPK pathway as acting in head cholinergic neurons to modulate *sax-7-*dependent locomotion. Loss of either KSR-1 or MPK/Erk dramatically suppresses the *rab-3; sax-7* and *unc-13; sax-7* abnormal locomotion, coiling tendency, and reduced swim and crawl rates (Fig 4, 5). Head cholinergic neurons comprise a subset of motor neurons that synapse on muscles in the head region as well as interneurons that synapse onto ventral nerve cord motor neurons to coordinate locomotion (White *et al.* 1986; Pereira *et al.* 2015). The MAPK pathway was previously discovered to also modulate locomotion mediated by the heterotrimeric G protein Gq (Coleman *et al.* 2018). Hyperactive Gq signaling results in aberrant loopy locomotion that is also dependent on KSR-1 function in head cholinergic neurons. Furthermore, an activated form of the MAPK component, LIN-45/Raf, expressed in the same head neurons in wild-type animals not only phenocopies this loopy posture but also results in aldicarb hypersensitivity. The MAPK signaling pathway may thus promote coordinated locomotion by modulating central patterning of synaptic activity.

The overlapping loopy posture of *rab-3; sax-7* and hyperactive Gq animals combined with the shared genetic interaction with the MAPK pathway suggest that SAX-7 may act to modulate the MAPK pathway specifically by dampening MAPK activity. In support of this notion, mammalian L1 is known to regulate MAPK signaling (Schmid *et al.* 2000; Cheng *et al.* 2005). It is conceivable that SAX-7 could influence MAPK activity either in the same cells or in adjacent cells with SAX-7 functioning in a non cell-autonomous fashion. Indeed, SAX-7 has been shown to influence the behavior of adjacent cells through physical interactions of its extracellular domain to diverse transmembrane proteins on adjacent cells (Sundararajan *et al.* 2019). In this study, we find that driving *sax-7* expression in head cholinergic neurons showed, at best, modest suppression of *rab-3; sax-7* uncoordinated locomotion (Fig 8), suggesting that *sax-7* does not act in the same neurons as *ksr-1*. Alternatively, it is possible that the poor rescue may be due to technical issues. For example, a recent study revealed that the strength of the *unc-17* promoter may be weak relative to other promoters for cholinergic interneuron expression (Bhardwaj *et al.* 2020). Additionally, somatic CRE activity may not be sufficient in our inducible knock-in system, resulting in insufficient SAX-7 levels in head cholinergic neurons for rescue of *rab-3; sax-7* phenotypes.

On the other hand, our results could point to a requirement for *sax-7* in additional neurons, besides head cholinergic neurons, raising an alternative or additional mechanism by which *sax-7* may coordinate locomotion. Previously, the mammalian L1CAM encoded by the CHL1 gene was shown to function as a co-chaperone, along with the 70 kDa heat shock protein (hsc70), the alpha cysteine string protein (αCSP), and the small glutamine-rich tetratricopeptide repeat-containing protein (αSGT), all chaperones of the exocytotic machinery that include the SNARE complex. CHL1 knockout mice exhibit reduced levels of assembled SNARE complex after stress or prolonged synaptic activity as well as impaired synaptic vesicle recycling, which became more apparent over time (Leshchyns’ka *et al.* 2006; Andreyeva *et al.* 2010). It is possible that *sax-7* functions in synapses in a similar manner. Such a role may account for why neuronal dysfunction in *sax-7* animals was apparent in the more rigorous swim assay and not with the crawl assay (Fig 2). Potential SV recycling defects in *rab-3; sax-7* neuromuscular junctions would likely not have been detected in our ultrastructural analyses and electrophysiological recordings because these animals were not subjected to prolonged synaptic stress prior to examination. The presynaptic chaperones and exocytotic machinery also function in the regulated exocytosis of dense core vesicles, which contain neuropeptides and hormones as opposed to classical neurotransmitters (Johnson *et al.* 2010; Burgoyne and Morgan 2015). Indeed, CHL1 was demonstrated more recently also to influence the translocation and/or docking of insulin-containing DCV vesicles to the plasma membrane of pancreatic islet cells, with loss of CHL1 leading to impaired insulin secretion. Furthermore, CHL1 is implicated as a risk factor in type 2 diabetes (Taneera *et al.* 2012; Xin *et al.* 2016; Jiang *et al.* 2020). It is thus also conceivable that SAX-7 could similarly influence exocytosis of neuropeptide-carrying DCVs. Neuropeptides play an important role in influencing *C. elegans* synaptic strength and behaviors (Li and Kim 2008). Intriguingly, loss of neuropeptides encoded by the *flp-1* gene results in changes in synaptic activity and a similar loopy body posture exhibited by *rab-3; sax-7* animals as well as by animals with elevated Gq signaling (Nelson *et al.* 1998; Stawicki *et al.* 2013; Buntschuh *et al.* 2018; Oranth *et al.* 2018).

To conclude, this study uncovered a role for the MAPK signaling pathway in SAX-7-dependent locomotion. While many questions remain regarding the mechanisms by which MAPK and SAX-7 coordinate locomotion and control body posture, the well-endowed genetic toolkit, mapped neural connectome, and well-characterized behaviors of *C. elegans* provides an excellent platform to further investigate the processes by which SAX-7 intersects the MAPK pathway.

## Acknowledgements

We thank Michael Ailion for the DNA constructs used to drive *ksr-1* expression in acetylcholine neurons and Guillermo Marques at the University of Minnesota Imaging Center for assistance with imaging and use of the NIS-Elements AR Analysis software used to quantify crawl rates. Some strains were provided by the *C. elegan*s Genetics Center, which is funded by the NIH Office of Research Infrastructure Programs (P40 OD010440). EN sample processing was partly performed at the BioCryo facility of Northerwester University’s NUANCE Center, which is supported by NSF (NSF ECCS-1542205 and NSF DMR-1720139) and the International Institute for Nanotechnology (IIN). EM images were acquired using instruments in the Electron Microscopy Core of the University of Illinois at Chicago’s Research Resources Center. This work was supported by NIH grant NS045873 to L.C.

---

**S1. Movies illustrating the synthetic Unc locomotion with coiling tendencies of *rab-3; sax-7* double mutants.**

In contrast to A) Wild-type, B) *sax-7*, or C) *rab-3* animals, D)*rab-3; sax-7* double mutant animals have synthetic locomotion behavior. These movies were captured on a Nikon AZ100 macroscope at 1 frame/second for 120 seconds and processed at 6 frames/ second with ImageJ (Schneider *et al.* 2012).

**Fig S2.**
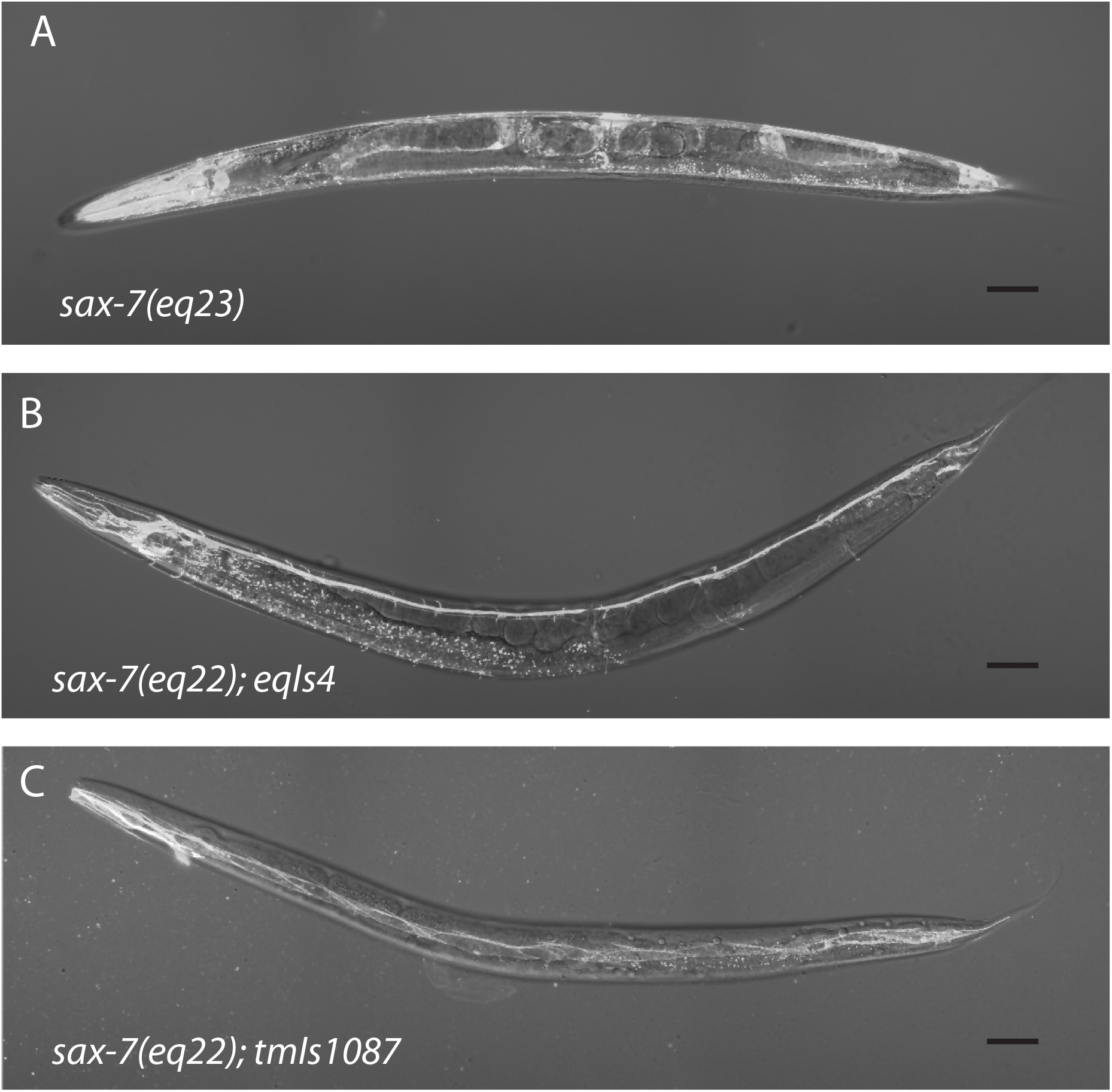
An overlay over DIC images of *sax-7*(*eq22*) animal showing restored SAX-7::mCherry using tissue-specific Cre-recombinase expression. (A) Cre-recombinase expression in the *sax-7(eq22)* germline results in the excision of the gene-disrupting cassette to generate *sax-7(eq23)* with endogenous SAX-7::mCherry restored in all tissues. A DIC image of a *sax-7(eq23)* adult animal with an overlay of its SAX-7::mCherry fluorescence signal. Crossing the P*_rab-3_*::Cre or P*_myo-3_*::Cre transgene, *eqIs4* and *tmIs1087* respectively, into the *sax-*7*(eq22)* strain results in somatic excision of the gene-disrupting cassette to restore endogenous SAX-7::mCherry expression in (B) neurons and (C) body-wall muscles, respectively. Scale bar 50 μm

**S3. Movies illustrating the suppression of *rab-3; sax-7* synthetic Unc locomotion in *ksr-1* and *mpk-1* null backgrounds.**

Both A) *rab-3; sax-7; ksr-1* and B) *rab-3; mpk-1; sax-7* animals show relatively wild-type locomotion. These movies were captured on a Nikon AZ100 macroscope at 1 frame/second for 120 seconds and processed at 6 frames/ second with ImageJ (Schneider *et al.* 2012).

**Fig S4.**
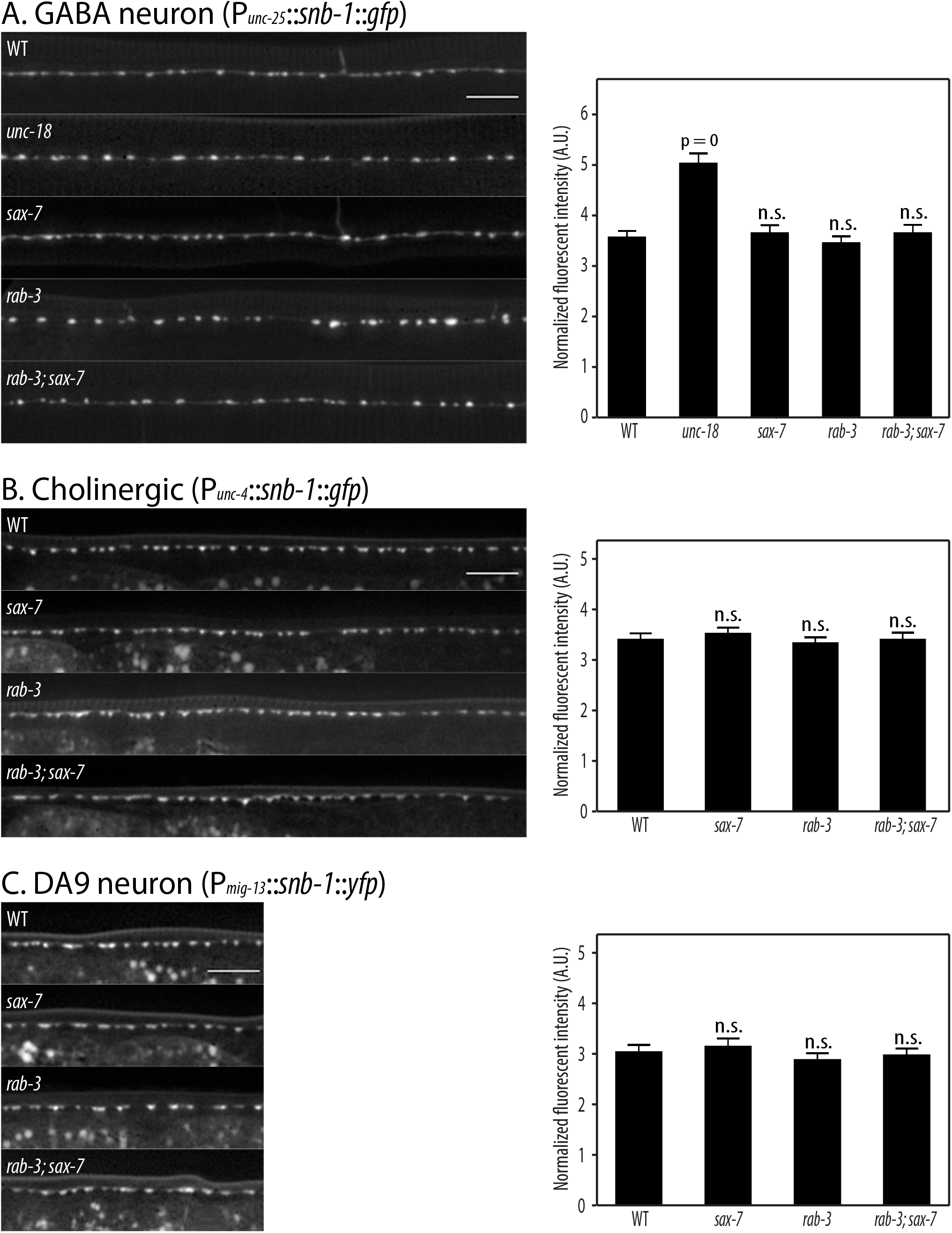
*In vivo* imaging and quantitation of the synaptic protein SNB-1::GFP reveals no apparent abnormalities in *rab-3; sax-7* synapses. *In vivo* imaging and quantitation of SNB-1::GFP reveal no apparent abnormalities in the number or intensity of *rab-3; sax-7* (A) GABA and cholinergic (B and C) synapses. In contrast, control *unc-18* animals (A) show the increased punctal intensity expected in animals with impaired synaptic vesicle release. Scale bar 10 μm

